# Ultrastructure of salivary glands in *Psammotettix alienus* Dahlbom (Homoptera: Cicadellidae) infected with Taastrup virus

**DOI:** 10.1101/263194

**Authors:** Thorben Lundsgaard

## Abstract

Taastrup virus (TV) is a novel virus belonging to *Mononegavirales* and with filovirus-like morphology. In adult *Psammotettix alienus* infected with TV, the highest concentration of virus particles was found in salivary glands, consisting of a principal gland (type I-VI-cells) and an accessory gland. Examination of thin sections revealed enveloped particles, about 1300 nm long and 62 nm in diameter, located singly or in paracrystalline arrays in canaliculi of type III- and IV-cells. In gland cells with TV particles in canaliculi, granular masses up to 15 micrometer in diameter are present in the cytoplasm. These masses are believed to be viroplasms, the sites for viral replication. TV particles were observed at the connection between a canaliculus and the salivary duct system.

## Introduction

Filovirus-like particles of Taastrup virus (TV) have been described from a population originating from France of apparently healthy leafhoppers belonging to the species *Psammotettix alienus* Dahlbom (Homoptera: Cicadellidae). The flexuous particles, 55-70 nm in diameter and 600 or 1100 nm long, consist of an inner coiled nucleocapsid about 30 nm in diameter and surrounded by a membrane with pronounced surface projections inserted (Lundsgaard, 1997), thus resembling members of *Filoviridae* morphologically. The RNA genome has partly been sequenced, which demonstrated both that TV belongs to the virus order *Mononegavirales*, but that it does not fit into any of the established families of this virus order (Bock *et al.*, 2004).

The hosts for *Psammotettix alienus* are grasses and cereals (Raatikainen and Vasarainen, 1976). The insects salivate during probing, penetration, and plant fluid ingestion (Backus, 1985) and it is thus likely that salivary glands play a key role in the epidemiology of TV. A detailed study on the ultrastructure of these organs from infected leafhoppers was initiated and the results presented here.

## Materials and Methods

The following experiments were performed in order to determine the distribution of virus in leafhoppers taken from a TV-infected population. Two fractions were made from ten randomly chosen insects. One fraction consisted of heads (including salivary glands) and the other fraction of thorax, abdomen, wings and legs. Each of the fractions was extracted in 1 ml distilled water. After one cycle of differential centrifugation (5 min at 1800 ×g and 20 min at 13,000 × g), the sediments were dissolved in 20 μl (head fraction) or 100 μl (body fraction) of negative staining solution (0.5% ammonium molybdate, pH 7). Preparations were made on Formvar^®^ mounted nickel grids (400-mesh) and the number of TV particles in ten grid squares was counted in a transmission electron microscope (JEOL JEM-100SX). The mean number of particles for the head fraction was 9.1 [standard deviation (s) = 4] and zero particles for the body fraction. In the next experiment, the heads from ten insects were severed and two fractions prepared, namely: a salivary gland fraction (SG) and a fraction representing the remaining part of the head (H). After differential centrifugation, both sediments were dissolved in 20 μL staining solution. The mean number of particles per grid square was 34.3 (s = 5) for SG and 0.3 (s = 0.7) for H, showing the salivary glands to be the most important site for accumulation of TV particles.

Ten adults taken in random from the TV infected population were dissected in Ringer’s solution (0.7% NaCl, 0.035% KCl, 0.0026% CaCL). Each pair of salivary glands was divided in a left and a right part with the purpose of checking one part for presence of particles by negative staining electron microscopy (extraction in 10 μl 0.5% ammonium molybdate) and the other corresponding part by ultrastructural analysis. The salivary glands were fixed, dehydrated, and embedded essentially according to Berryman and Rodewald (1990). In brief, the specimens were transferred to fixative (3% formaldehyde, 0.3% glutaraldehyde, 100 mM phosphate buffer, pH 7.0) and maintained in fixative for at least 2 h. After washing (3.5% sucrose, 0.5 mM CaCl_2_, 100 mM phosphate buffer, pH 7.4), the aldehyde groups were quenched with 50 mM NH_4_Cl dissolved in washing solution) for 1 h. The specimens were then washed (3.5% sucrose, 100 mM maleate buffer, pH 6.5) and post-fixed with 2% uranyl acetate in the same maleate buffer for 2 h. After dehydration in acetone (50%, 70%, 90%), the specimens were infiltrated in LR Gold (The London Resin Co., England) containing 0.5% benzoin methyl ether (3 changes) and polymerised under near ultraviolet light (Philips TW6). Fixation and dehydration (including 50% acetone) was done at 5-10 °C and dehydration (from 70% acetone), embedding, and polymerisation performed at −20°C. Thin sections (60-70 nm) were collected on 200-mesh nickel grids, stained with saturated uranyl acetate for 5 min and lead citrate (1 mM, pH 12) for 1 min, and examined with a JEOL 100SX transmission electron microscope operated at 60 kV. As controls, specimens were prepared in the same way from a healthy population of *P. alienus.*

## Results and Discussion

Sogowa (1965) has described the morphology of leafhopper salivary glands in detail, but *P. alienus* was not included and has not been described elsewhere. The morphology of these glands was therefore studied and described here. The naming of the glandular cells made by Sogowa (1965) is followed. The salivary glands of P. *alienus* are lying in the head and prothorax and consist of two pairs of glands, each made up of a complex principal gland (Fig. 1, I-VI) and an U-formed accessory gland (Fig. 1, AG). The lettering III, IV, and V all represents single cells, but the areas designated I, II, and VI in Fig. 1 represents one to more cells each (the exact number not determined). The accessory salivary gland (AG) consists of many cells. According to the ultrastructure of the canaliculi in type III-cells (see later), I have subdivided these cells in four type IIIa-cells and two type IIIb-cells. The orientation of the principal salivary gland within the insect is by having the one IIIb-cell pointing downward and the other IIIb-cell against the front. The I- and II-cells (anterior lobe) form the proximal part and the IV-through VI-cells (posterior lobe) the distal part of the gland. The overall length of a typical principal salivary gland (from the tip of the lower IIIb-cell to the tip of the upper IIIa-cell) is about 550 μm. Negative staining electron microscopy revealed TV particles in glands from two of ten examined insects. By ultrastructural analysis, unusual structures, as described below, were detected in glands from the same two individuals. Rod-shaped particles were observed either singly or in paracrystalline arrays in canaliculi of type III- and IV-cells (Fig. 2–4). The type III-cells are easily distinguished from other salivary gland cells by not having secretory vesicles in the cytoplasm. However, two types of III-cells could clearly be distinguished in *P. alienus* according to the structure of canaliculi. In the one type, here designated IIIa, the content of the canaliculi is granular (Fig. 1 and 2), whereas the other, designated IIIb, has canaliculi filled with many small membrane-bound spherules containing electron-opaque granules (Fig. 1 and 3). The IV-cells could easily be identified from all other salivary gland cells by a visible line pattern in their secretory vesicles (Fig. 4).

**Fig. 1.**
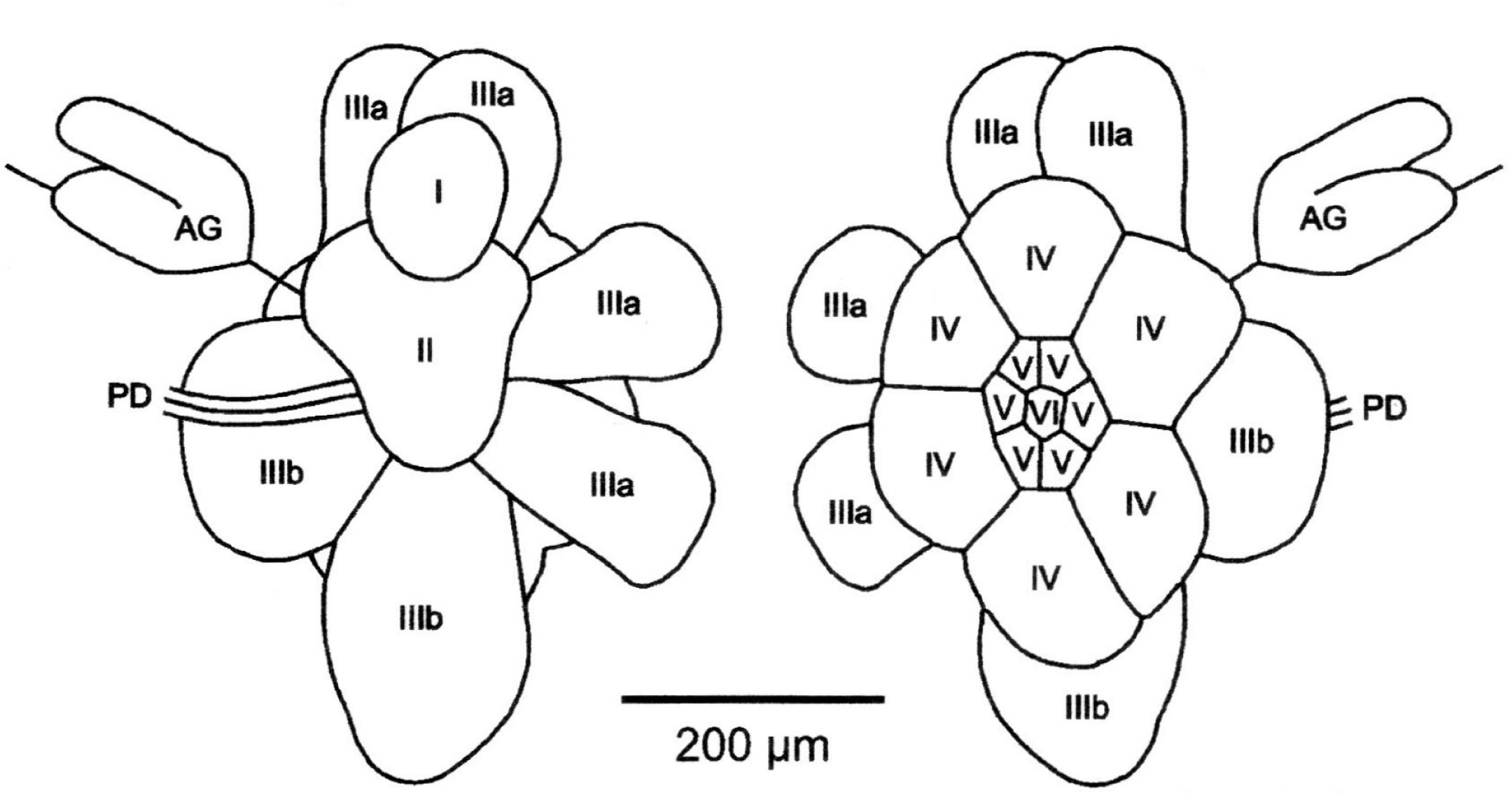
Drawing of salivary glands from *Psammotettix alienus* seen from the insect axis (left) or from outside (right). The salivary glands consist of a principal gland (cell type I to VI) and an accessory gland (AG), both connected to the principal duct (PD). According to ultrastructure of canaliculi, the type III-cells are differentiated in four type IIIa-cells (pointing upwards and backwards) and two type IIIb-cells (pointing forwards and downwards).

**Fig. 2.**
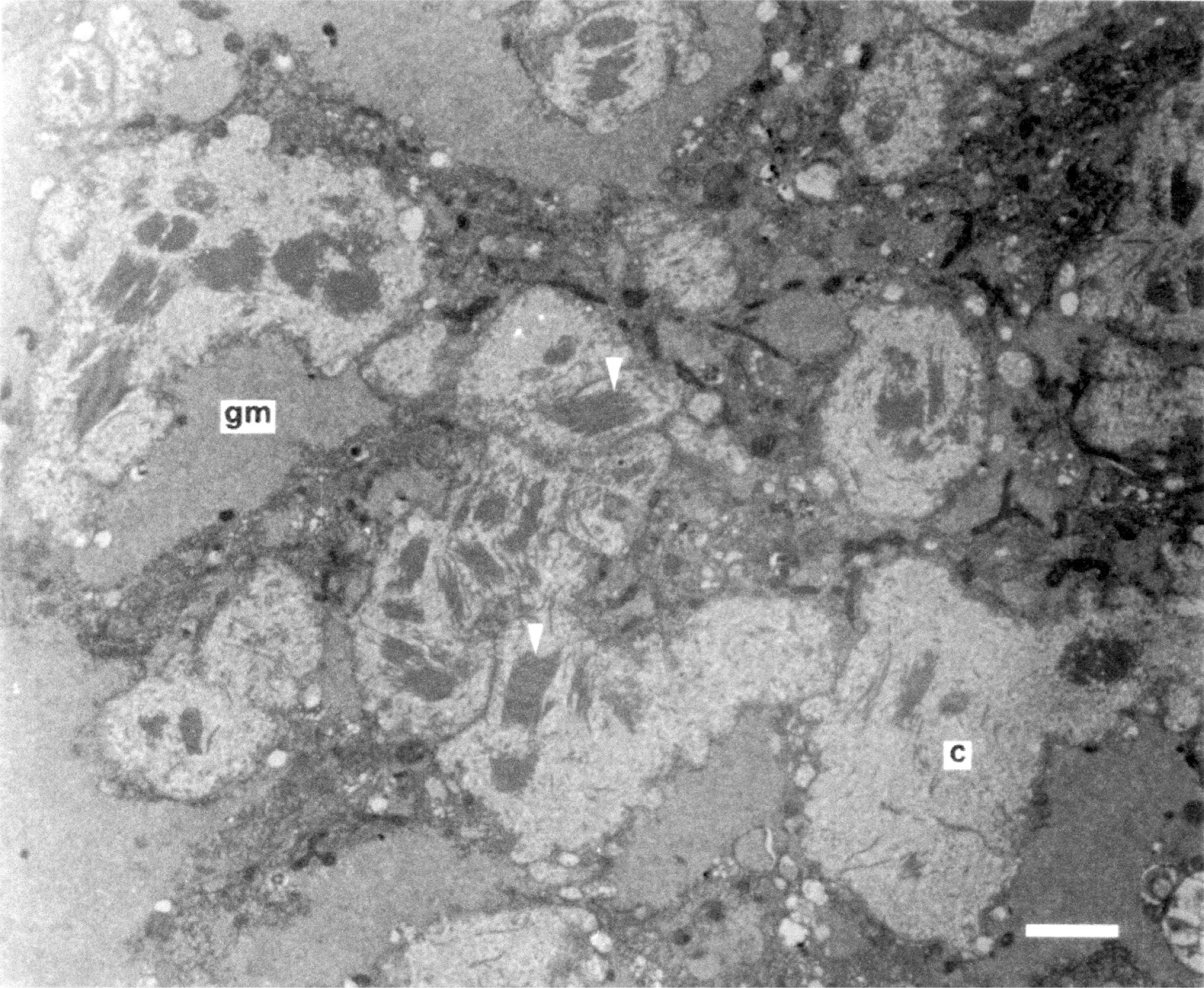
Section through a type IIIa-cell from *P. alienus* infected with Taastrup virus. Aggregates of virus particles (white arrowheads) are located in canaliculi **(c)** close to granular masses **(gm)** in the cytoplasm. Bar = 2 μm.

**Fig. 3.**
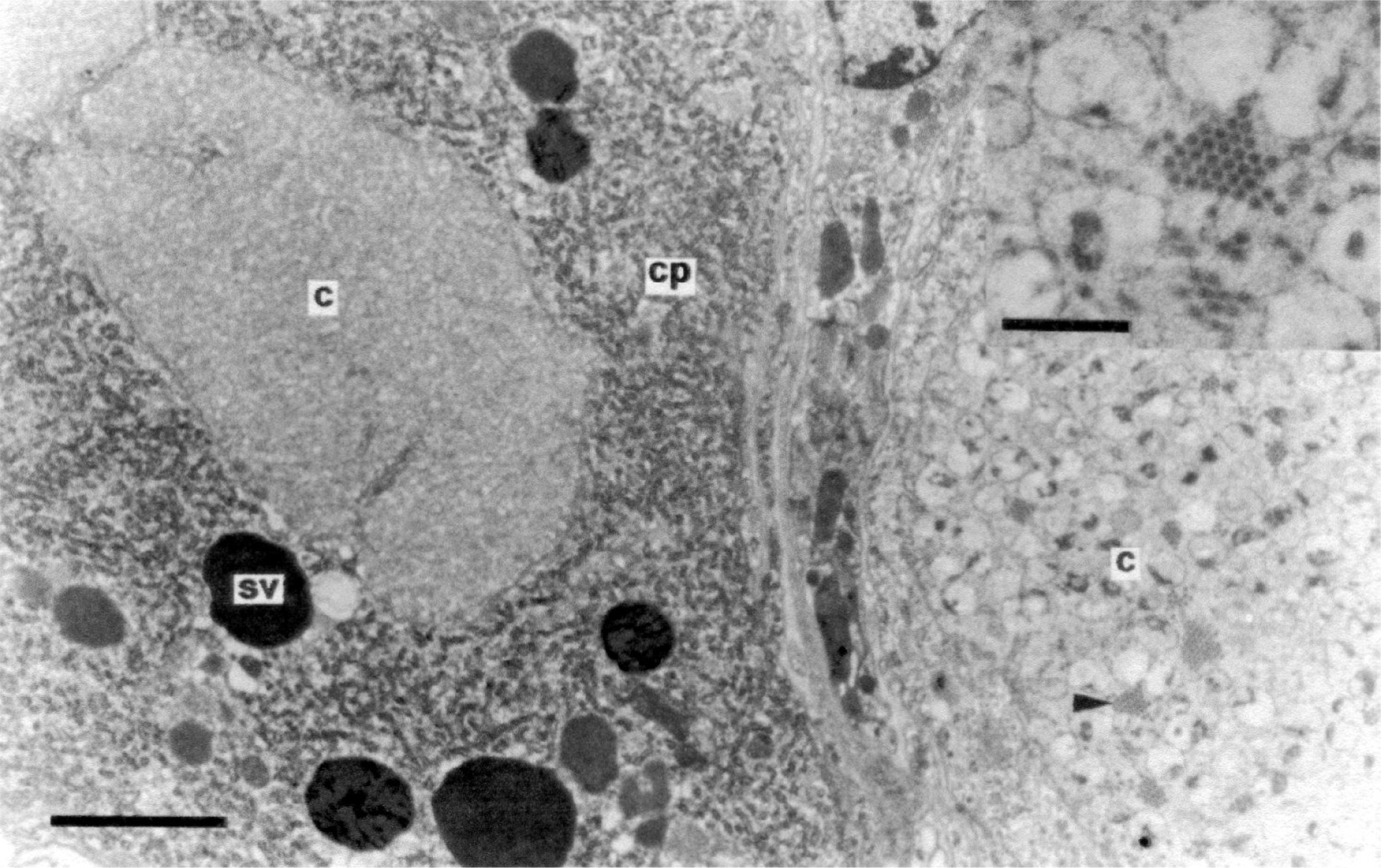
Section through a type IIIb-(right) and a type IV-cell (left) from *P. alienus* infected with Taastrup virus. Aggregates of virus particles (black arrowhead) are located in a canaliculus of the type IIIb-cell (inset). Note the vesicles with electron opaque granules in the canaliculus, characteristic of type IIIb-cells. The neighboring cell (type IV) has cytoplasm **(cp)** with rough endoplasmic reticulum together with electron opaque secretory vesicles (sv). No virus particles were observed in any of the canaliculi (c) of this cell. Bar = 2 μm. Inset bar = 500 nm

**Fig. 4.**
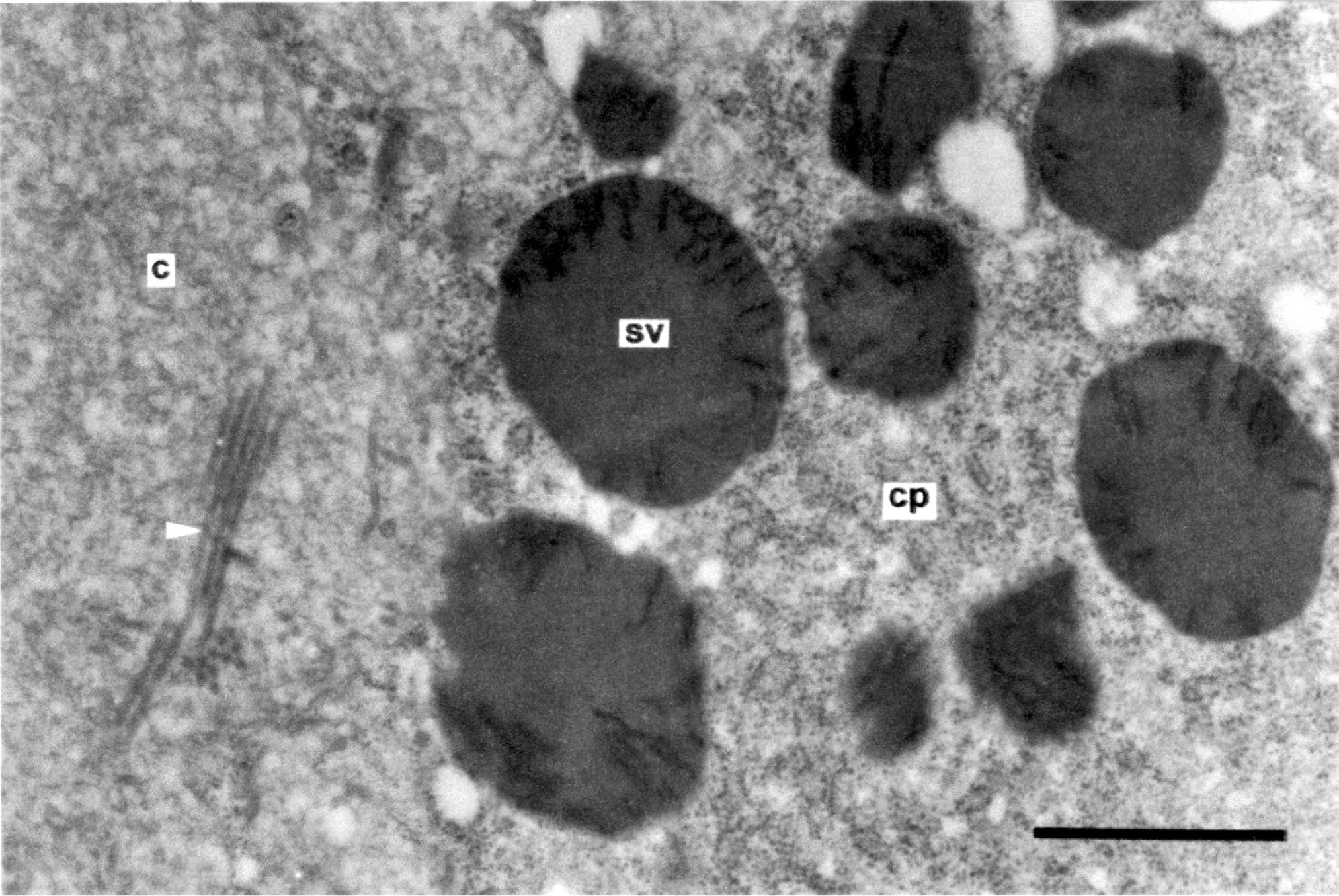
Section through a type IV-cell from another infected leafhopper than that examined in Fig. 3. Virus particles (white arrowhead) are located in canaliculi **(c)** of this cell. Note the dark lines in the secretory vesicles **(sv)**, which are characteristic for the type IV-cells. **cp** = cytoplasm. Bar = 1 μm.

The particles in the canaliculi were often seen with a membrane-like bleb at the one end (Fig. 5). These blebs are probably remnants of viral envelopes remaining after a presumed budding process. For particles for which both ends could be seen, the mean length - not including the bleb - was calculated to be 1294 nm (s = 22). In cross section, an electron-opaque inner hollow core (about 31 nm in diameter) is seen surrounded by an outer faint electron opaque layer, separated by a translucent layer (Fig. 6). The diameter of particles was calculated to be 62 nm from particles laying in paracrystalline arrays. The morphology and dimensions for the particles observed here in canaliculi are in agreement with the negative stained particles presented previously (Lundsgaard, 1997). This taken together with the correlation found between presence of particles in extracts and presence of particles in canaliculi among the ten examined leafhoppers, I conclude that the particles observed in canaliculi are identical to those described in the former paper. So, the inner, hollow core is probably the helical nucleocapsid of TV and the outermost faint layer the G protein spikes. In cross sections through particles of rhabdoviruses and filoviruses (Murphy and Harrison, 1979; Geisbert and Jahrling, 1995), an electron opaque layer, believed to be the virus membrane, is present between the layer of spikes and the inner nucleocapsid core. The analogous position of this electron opaque layer is translucent in the present TV particles. This discrepancy can be explained by use of osmium tetroxide for postfixation (known to preserve and stain membranes) in the studies on rhabdo- and filoviruses, whereas osmium tetroxide was omitted in the present investigation. Osmium tetroxide was omitted in order to preserve the antigenic activity for future research on localisation of viral antigens in these specimens. Aggregates of particles, 75-80 nm in diameter, have been described from head cells of *P. alienus* (Lundsgaard, 1997). These particles were seen located in the cytoplasm (not in canaliculi as presented here) and their central, electron opaque core was not seen hollow (as are the present TV particles), suggesting that the particles described previously were not TV particles.

**Fig. 5.**
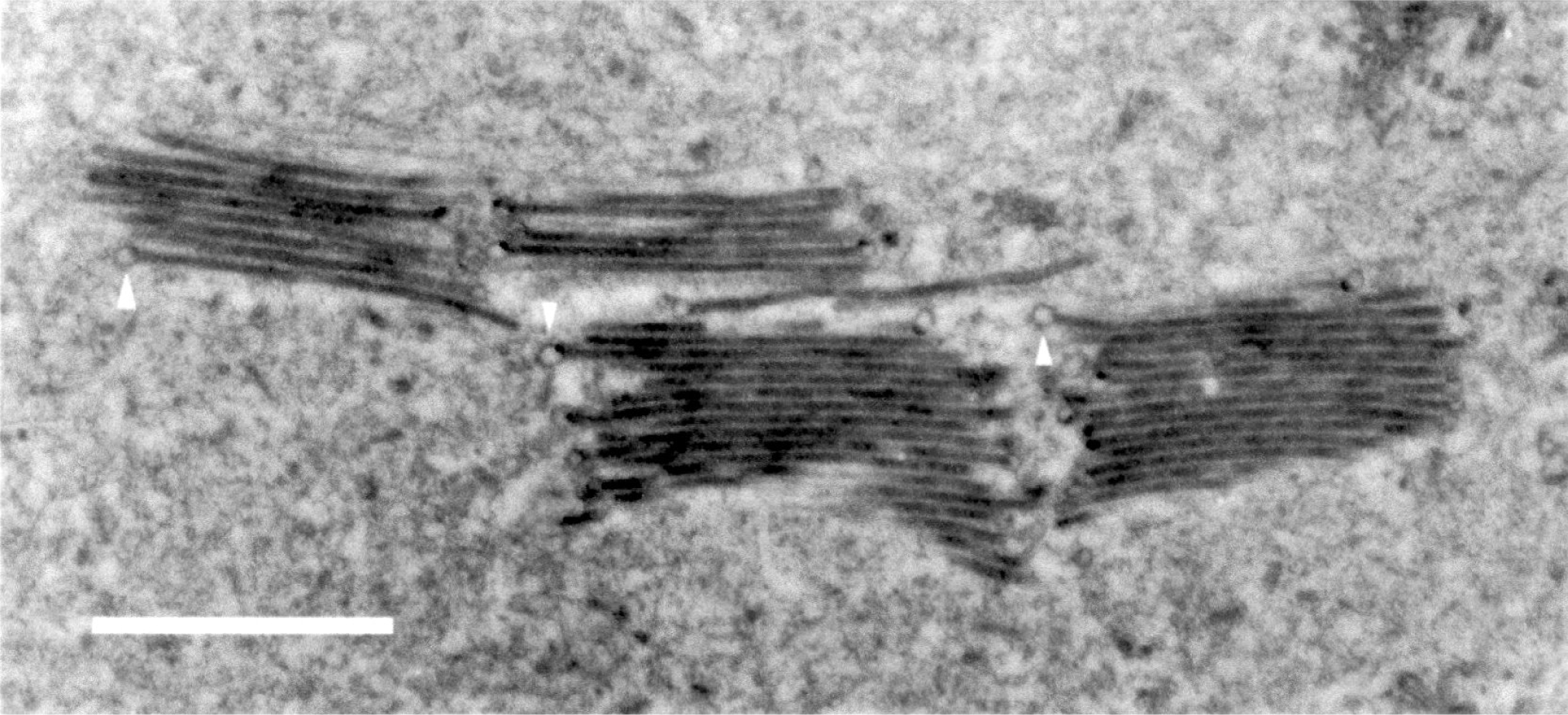
Longitudinal section through aggregates of virus particles located in a canaliculus of a type IIIa-cell from *P. alienus* infected with Taastrup virus. Small blebs (white arrowheads) are seen at one end of several particles. Bar 1 μm.

**Fig. 6.**
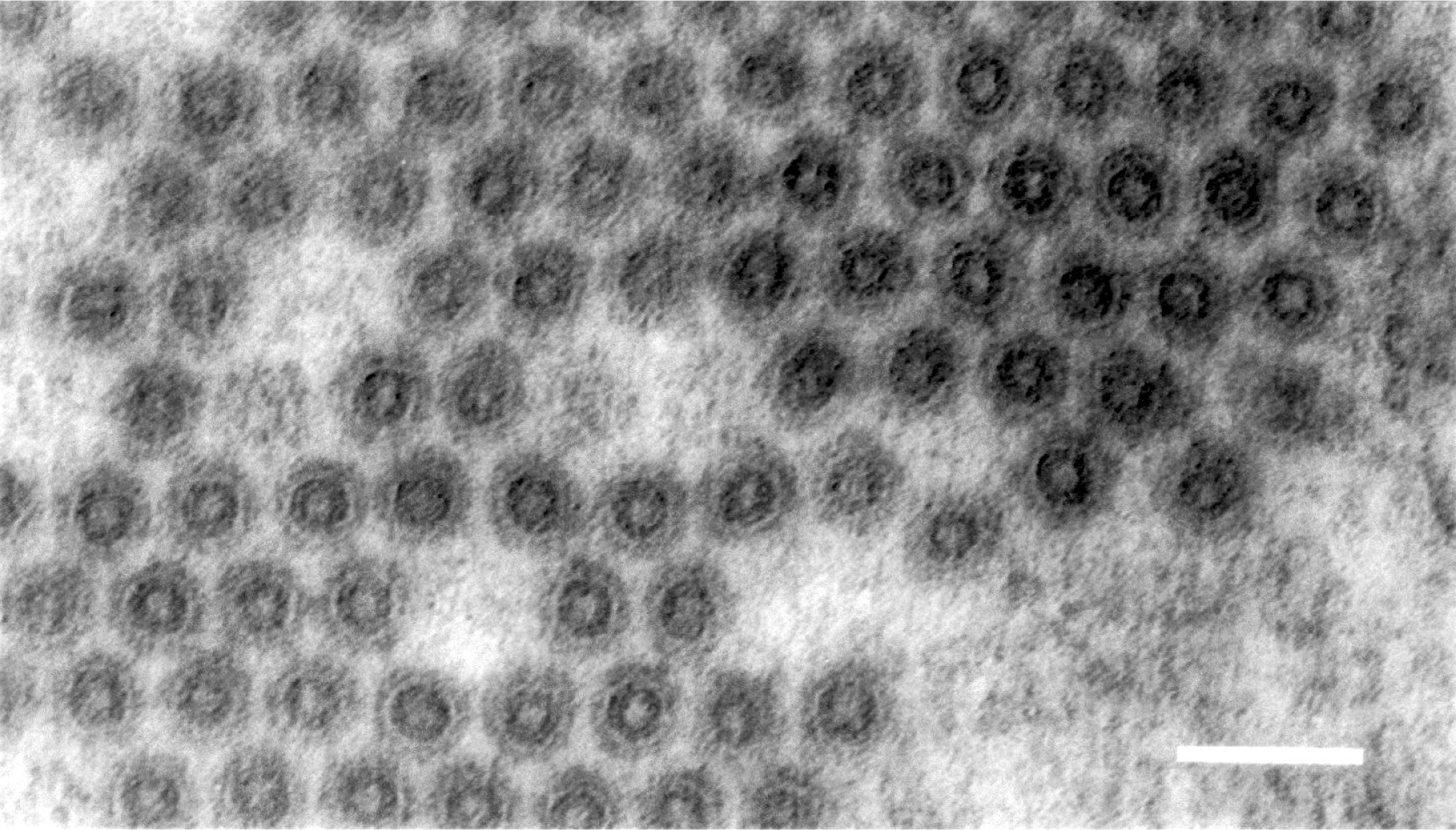
Transverse section through virus particles located in a canaliculus of a type IIIb-cell from *P. alienus* infected with Taastrup virus. An electron opaque tube with a hollow centre is seen surrounded by a translucent layer and outermost a faint electron opaque layer. Bar = 100 nm.

In the cytoplasm of type III- and IV-cells, possessing canaliculi with TV particles, fine granular masses up to 15 μm in diameter were observed (Fig. 2). These masses do not contain ribosomes or other cellular organelles and their presence was positive correlated with presence of TV particles in nearby canaliculi. Granular masses (viroplasms), believed to be viral replication sites, have been observed in cells infected with rhabdoviruses (Murphy and Harrison, 1979; Conti and Plumb, 1977) and filoviruses (Geisbert and Jahrling, 1995). The granular masses present in the cytoplasm of salivary glands with TV particles are thus suggested to be viroplasms, the sites for TV replication. Because a nuclear localisation signal has been identified in the G-protein gene of TV (Bock *et al.*, 2004), nuclei close to viroplasms were carefully examined for abnormalities as compared with uninfected controls. Neither budding through nuclear membranes nor abnormality of nucleoplasm was seen during the present investigation.

In several cases, two glandular cells in contact with each other could be found by which the one contained viroplasms in the cytoplasm and did contain TV particles in canaliculi, while the other neighbouring cell was uninfected (Fig. 3). These uninfected cells were of a type, which was seen infected elsewhere. TV particles were never seen between such cells or in the space between the plasma membrane and the basal lamina, so, the delivery of mature TV particles seem only to take place by budding of nucleocapsids - synthesised in viroplasms - into canaliculi destined for saliva excretion. Figure 7 shows an oblique section through the outlet from a type IIIb-cell canaliculus into the salivary duct system. Several TV particles can be seen at this outlet (Fig. 8). Muscle tissue, in form of myofibrils, is located around this outlet together with a neuron cell (Fig. 7); both probably involved in controlling the saliva outlet from this cell.

**Fig. 7.**
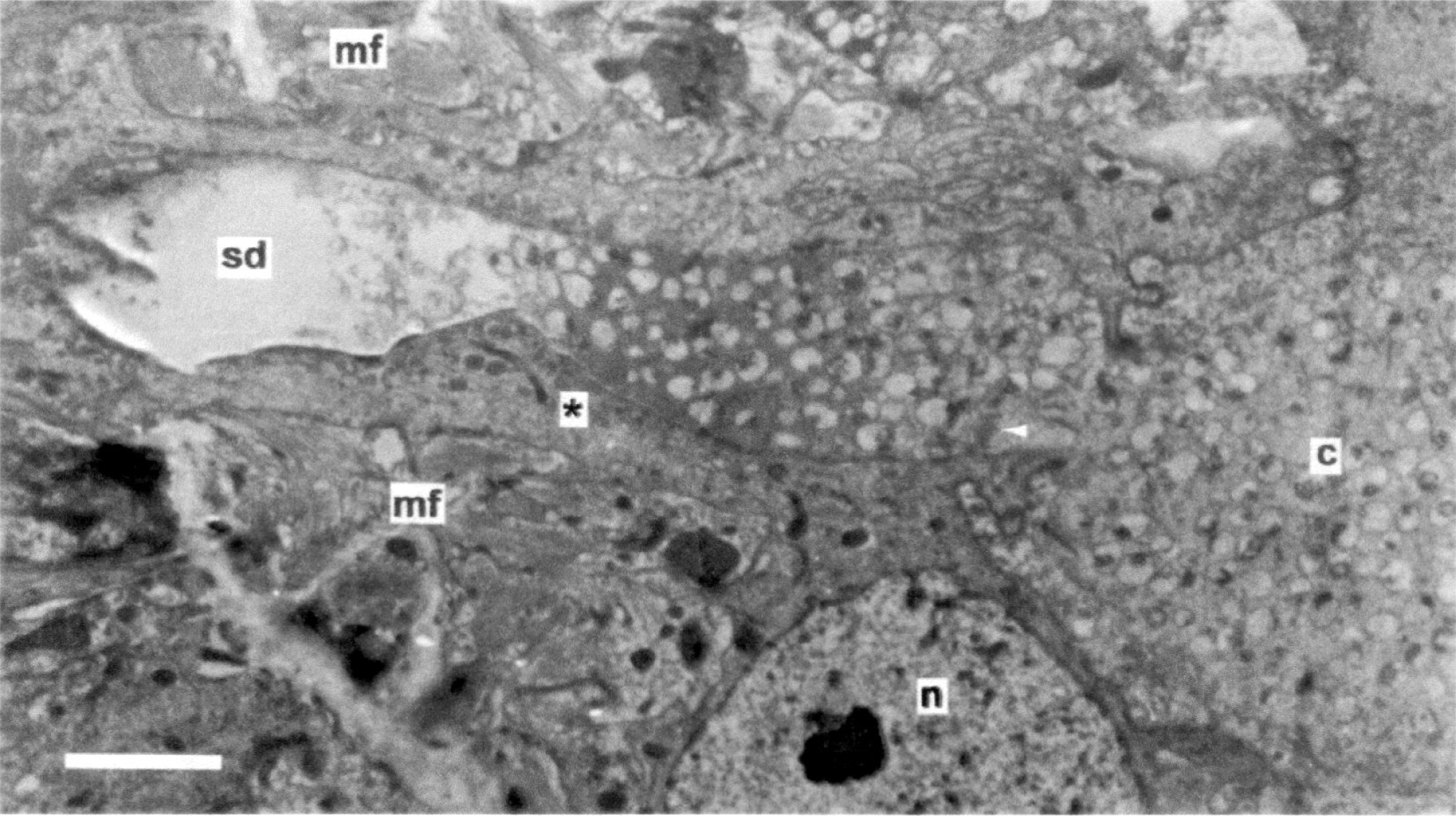
Oblique section through an outlet from a type IIIb-cell to the salivary duct system. The content of a canaliculus **(c)**, including virus particles (white arrowhead), is seen released into a salivary duct **(sd)**. Tissue with many myofibrils **(mf)** is located close to the salivary duct. A neuron with its nucleus **(n)** is seen with its axonic or dendritic extension (*) between the myofibrils and the salivary duct. Bar = 2 μm.

**Fig. 8.**
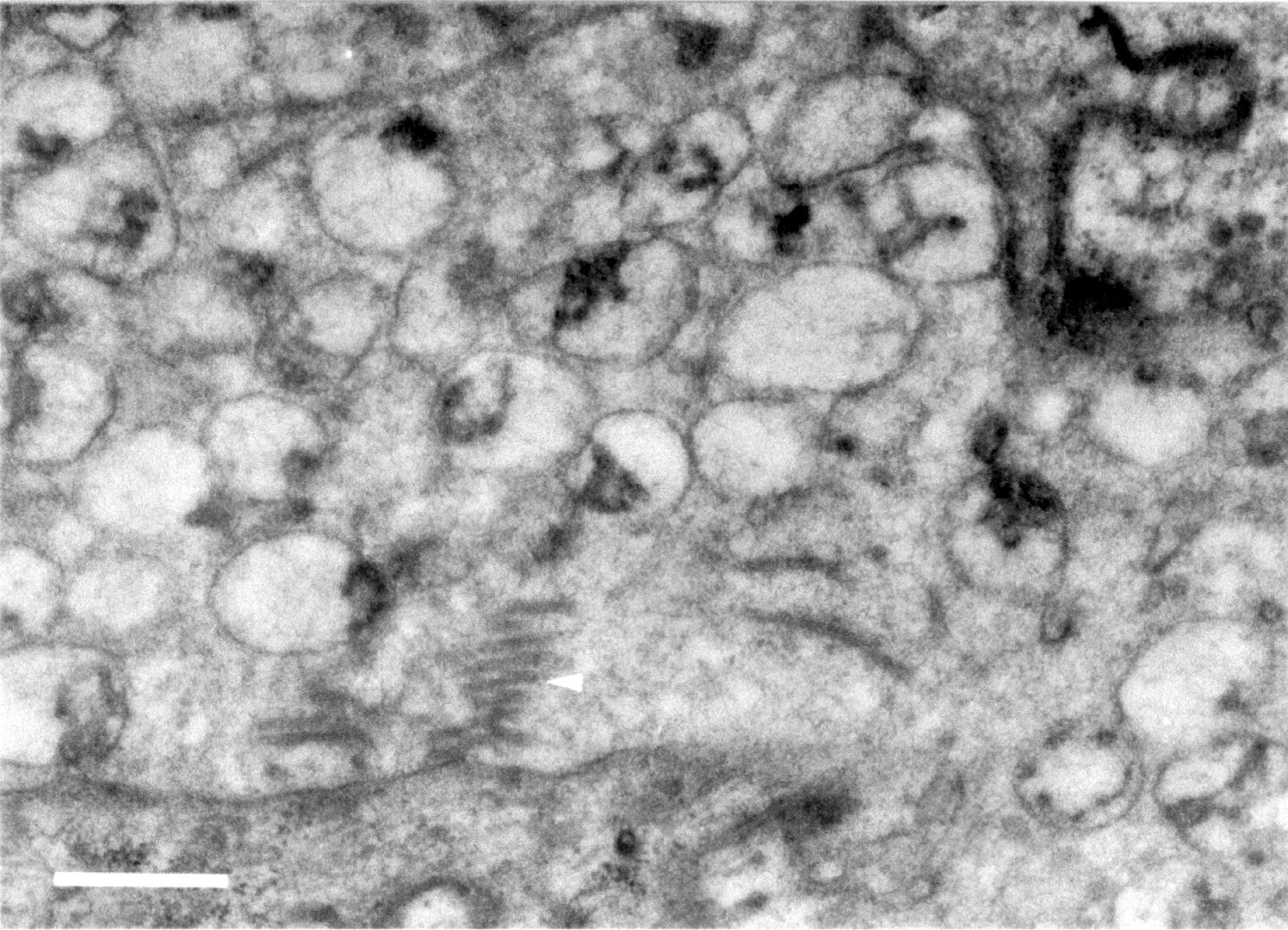
Close-up of the area in Fig. 7 where a canaliculus opens into the salivary duct system. Oblique sections through some virus particles are seen (white arrowhead). Bar = 500 nm.

TV-particles or viroplasms were neither observed in type I-, II-, V-, and VI-cells of principal salivary glands, nor in accessory salivary glands from infected leafhoppers. TV-particles or viroplasms were not seen in salivary glands from leafhoppers originating from a TV-free population of *P. alienus.*

The present investigation shows that salivary glands are the primary site for TV synthesis in adult P. *alienus.* For making sense for the virus life cycle, the TV particles excreted with saliva into the plant tissue must either be inoculum for TV infection of the plant, or be passively delivered to the plant and then later taken up by young healthy nymphs. Former investigations (Lundsgaard, 1997) have shown, that host plants used for rearing TV infected leafhoppers, do not become infected, and so, the plants probably function as passive vectors for transmission of TV between leafhoppers. The expected early events of virus uptake and infection in the alimentary canal system of nymphs would be an interesting topic for future research.

